# Successive failure triggered by motor exploration in a reinforcement-based reaching task

**DOI:** 10.1101/2025.02.03.634095

**Authors:** Risa Saito, Mitsuaki Takemi, Midori Kodama, Toyo Ogasawara, Daichi Nozaki

## Abstract

Motor exploration, a key process in reinforcement-based motor learning, is triggered by suboptimal performance and leads to increased movement variability. Consequently, failure in one trial may increase the likelihood of failure in subsequent trials. If this pattern emerges, successive failures (SFs) are not merely events to be avoided but part of the trial-and-error process of identifying optimal movements, which can facilitate motor learning. This study investigated whether and how SFs occur above chance levels during a motor learning task with binary success/failure feedback and explored the relationship between SFs and motor learning outcomes. Thirty-three healthy young adults participated in the study, performing a reaching task to pass through a target without visual feedback of their hand position. Binary feedback was provided after each trial, indicating whether the hand trajectory overlapped with the target. The results showed that the probability of two SFs was significantly higher than the square of the overall failure probability, indicating that failure streaks occurred above chance levels. A computational model, in which motor variability comprised constant motor noise and exploratory variability regulated by recent reward history (where variability increased following failure), successfully explained SF emergence. However, the learning index, defined as the difference in failure probability between the first and latter halves of the experiment, was unrelated to SF emergence. Additionally, failure streaks disappeared when participants were asked to reach one of seven randomly selected targets in each trial, suggesting that introducing environmental variability can help alleviate SFs.

## Introduction

A widespread belief in the sports world holds that success breeds success and failure breeds failure, often encapsulated in the concept of a “hot hand” or “streak.” This concept describes periods when an athlete’s performance markedly exceeds expectations based on their overall performance (Bar-Eli et al., 2006). The “myth of the hot streak” has been debated for decades in sports psychology and sports economics. While the classic work by Gilovich et al. (1985) argued that the hot streak is a fallacy of the human mind, others have suggested that it is inappropriate to search for streaks in rich contexts where other effects can mask the phenomenon (Kaplan, 1990; Koehler et al., 2003). Miller and Sanjurjo (2021) provided robust statistical evidence of shooting streaks by focusing their study of the “hot hand” phenomenon on controlled settings like the NBA three-point contest.

Although statistical evidence exists, the neural underpinnings of the hot streak remain unclear. Previous neuroscience studies have suggested that repeating the exact same movement is virtually impossible due to inherent movement variability. This variability may result from a noisy nervous system (Cohen et al., 2009; Faisal et al., 2008; Renart et al., 2014; Stein et al., 2005) and thus impede motor performance (e.g., a series of successful three-point shots). According to this view, movement variability, an inevitable result of nervous system noise, acts as an error-inducing factor that impedes hot streaks rather than supporting their occurrence.

However, recent studies have advanced a complementary view of motor variability (MV) (Wu et al., 2014; Dhawale et al., 2017; Uehara et al., 2019). In motor learning, we acquire both an action-value map to select highly rewarded actions and an internal representation of novel dynamics to reduce sensory prediction errors. In controlled experimental environments, such as visuomotor rotation and velocity-dependent force field, participants are asked with reducing systematic errors, making the learning process primarily dependent on minimizing sensory prediction errors. Conversely, in practical skill learning, typically devoid of perturbations, performance improves by selecting highly rewarded actions and may rely more on success-based exploration than on adaptation. In this context, MV is considered an unwanted byproduct of a noisy nervous system (Churchland et al., 2006) but is also seen as a crucial factor for reinforcement motor learning. MV is modulated as a function of past rewards to guide new task solutions through exploration (Wu et al., 2014; Dhawale et al., 2017; Uehara et al., 2019).

Motor exploration, a crucial component of reinforcement motor learning, is triggered by poor performance, leading to increased movement variability (Pekny et al., 2015). Thus, failure in one trial may increase the likelihood of failure in subsequent trials. The magnitude of motor exploration is related to learning speed in reinforcement-based arm-reaching tasks that require movement modification toward an optimal pattern (Chen et al., 2017; Therrien et al., 2016; Wu et al., 2014). These findings suggest that successive failures (SFs) are not merely events to avoid but part of the trial-and-error process of determining optimal movements and can facilitate motor learning.

In this study, we investigated whether SFs occur above chance levels during a motor task with binary success/failure feedback and examined their relationship to motor learning outcomes. Understanding how the brain regulates MV as a function of performance history and task uncertainty is inherently challenging. Measuring how MV fluctuates over many trials as a function of context and task parameters (e.g., reward rate) requires a large volume of data (Dhawale et al., 2017). Despite evidence that changes in reward rates modulate MV (Pekny et al., 2015), the specific algorithm through which the human brain adjusts variability based on past performance remains unknown. Furthermore, whether MV fluctuations represent active exploration processes at optimizing performance or merely unwanted byproducts of nervous system noise remains unclarified (Therrien et al., 2018). To address this, we employed a computational model that accounts for both constant motor noise and exploratory variability, dynamically regulated as a function of recent reward history, to explain how the human brain regulates these two MV types. Finally, we examined the link between internal model parameters representing constant and adaptively regulated MV and the degree of motor performance improvements throughout the experiment.

## Results

### Successive successes and failures occurred above the chance level

In Experiment 1, we examined whether SFs during a motor task occur more frequently than by chance. This was tested using a task with reward-based feedback (binary success/failure). Thirty-three healthy young adults performed a reaching task in which they moved their right hand toward a target without visual feedback. Participants controlled a cursor displayed on a screen using a robotic manipulandum and were asked to move the cursor through a target presented 10 cm away from the starting position (Fig. 1a). Success or failure was determined based on whether the hand trajectory overlapped the target.

**Figure 1.**
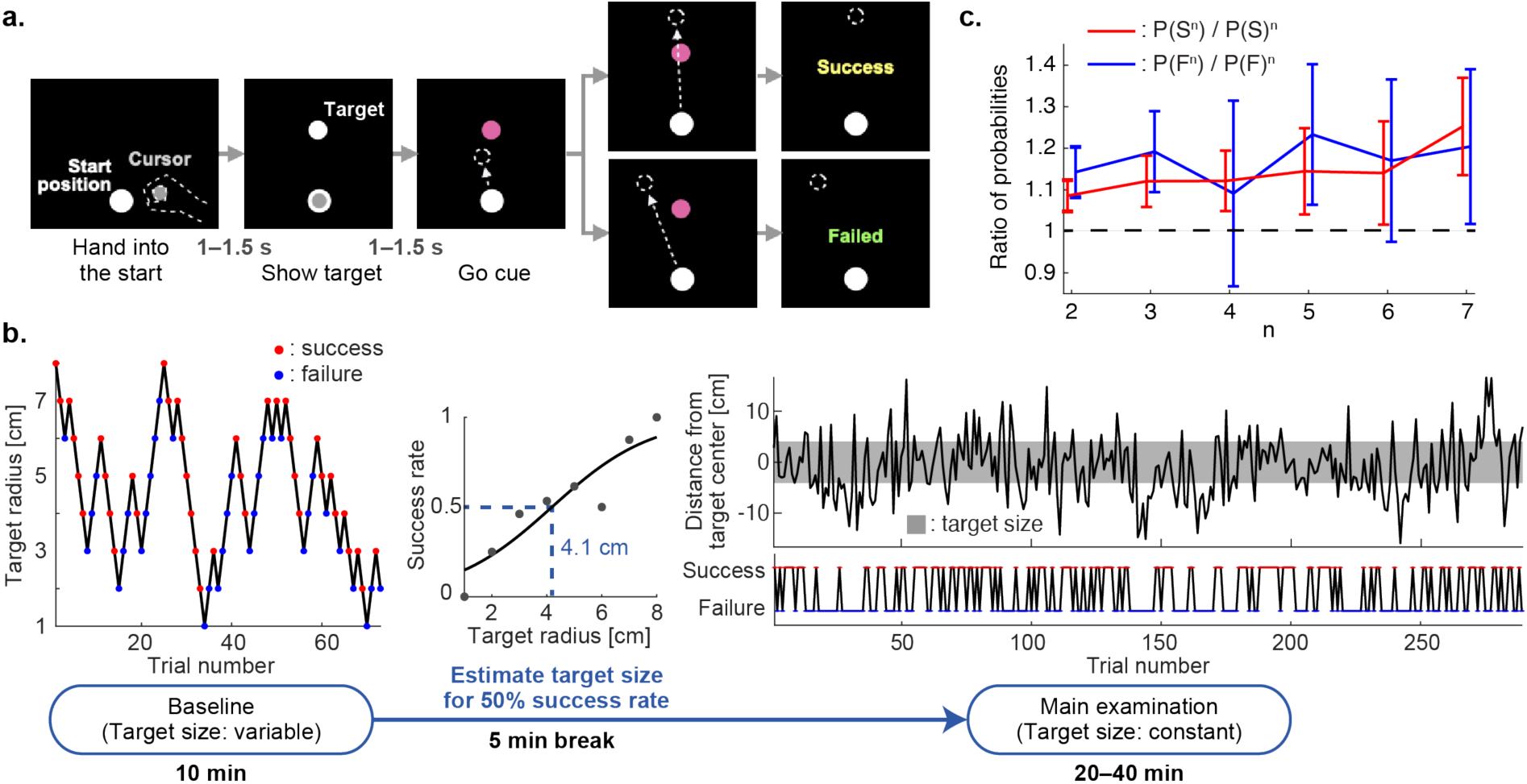
Success and failure streaks exceeded chance levels in a reinforcement-based motor task. (a) Illustration of the reinforcement-based motor task, wherein participants used their right hand to reach a target without visual feedback of hand position. Success or failure was determined based on whether the hand path overlapped with the target. (b) Experimental procedure. A baseline examination was conducted to estimate the individual target size corresponding to a 50% success rate. The scatter plot on the left shows individual trial data for target radius and success or failure. The middle plot depicts success rates for each target radius used to estimate the target size through sigmoidal curve fitting. The estimated target size was then used for the main examination. In the right plot, the gray shaded area represents the estimated target size for a 50% success rate. (c) Assessment of successive successes and failures. The red line represents the ratio of the probability of *n* successive successes P(S^n^) to the *n*-th power of the success probability for the main examination P(S)^n^. The blue line represents the ratio of the probability of *n* successive failures P(F^n^) to the *n*-th power of the failure probability for the main examination P(F)^n^. Data are represented as the mean ± 95% confidence interval. The lower boundary of the error bars exceeding 1 (indicated by the black dotted line) shows that the probabilities of the occurrence of *n* successive successes and failures significantly exceeded the chance levels.

Initially, participants engaged in a baseline task where success resulted in a 1-mm decrease in the target radius for the subsequent trial, while failure led to a 1-mm increase in the target size (Fig. 1b). The baseline task results were used to individually estimate the target size, which would yield a 50% success rate. The maximum, minimum, and average target sizes were 6.6, 2.1, and 4.2 cm, respectively. This individually adjusted target size was then used for the main examination, during which individual success rates were 33.8%–80.8%. The respective numbers of successive successes and failures were 4–34 and 4–29. Two-thirds of the participants showed longer than nine successive successes and six SFs.

We then calculated the probability of making a success in the current trial *n* given *n*−1 consecutive prior successes P(S_n_ | S_n-1_) for each participant (Fig. S1a). If the successive success is independent of prior successes, P(S_n_ | S_n-1_) = P(S), the overall success rate for the main examination. In other words, 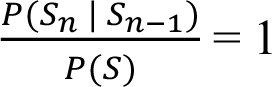. For 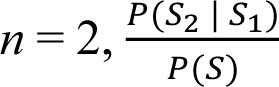 can be rewritten as 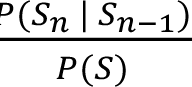, where P(SS) denotes the probability of making two successive successes. Thus, 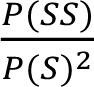 can be expressed as 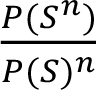, where P(S^n^) is the probability of making *n* successive successes.

The results showed that 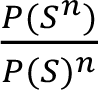 exceeded 1 for *n* = 2–7 (Fig. 1c, red line), indicating that successive successes consistently occurred above the chance level. For instance, the observed frequency of two successive successes was 1.086 ± 0.038 (mean ± 95% confidence interval (95%CI)) times higher than the square of the overall success rate. Similarly, we calculated the probability of making a failure in the current trial *n* given *n*−1 consecutive prior failures P(F_n_ |) for each participant (Fig. S1b) and found that 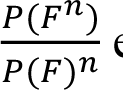 exceeded 1 at *n* = 2, 3, 5, and 7 (Fig. 1c, blue line). For example, the probability of two SFs was 1.143 ± 0.061 times higher than the square of the overall failure rate. Correlations between 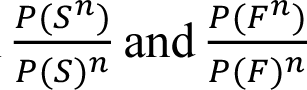 were significant for n = 2 (Fig. S1c left; r(31) = 0.503, p = 0.003) but not for n > 2. For instance, the results of n = 3 and 4 were r(31) = 0.138 (p = 0.44; Fig. S1c middle) and r(30) = 0.301 (p = 0.094; Fig. S1c right), respectively.

### Reward prediction regulated MV explains the emergence of successive successes and failures

Why do successive successes or failures occur? The reinforcement learning mechanism posits that failure triggers exploration, increasing movement variability, while success triggers exploitation, reducing movement variability. This mechanism may regulate the occurrence of successive successes and failures. To investigate this, we employed a computational model proposed in a previous study (Dhawale et al., 2019). In this model, the motor output direction *x* comprises three components (Fig. 2a): policy *µ* representing the intended movement direction, exploration *ε_e_* representing variability in movement for exploring new strategies, and motor noise *ε_n_* representing random errors in movement. The policy for the next trial *µ*(*i* + 1) is updated based on the learning rate *α_µ_*, reward prediction error at the current trial *δ*(*i*), and exploration at the current trial *ε_e_*(*i*). *ε_e_* is normally distributed with exploratory variability *σ_e_^2^*, which is updated in each trial depending on the amount of reward prediction *r̂* (Fig. 2b). Thus, the more the participants fail the task, the lower the reward prediction, the wider the exploration, and the greater the variability of the movement direction. *ε_n_* is normally distributed with variance *σ_n_^2^*, which remains constant throughout the experiment. The equation *σ_e_^2^* = *λr̂*(*i*) + *c* suggests that exploration variability is a linear function of reward prediction (Fig. 2c), where *λ* and *c* are free parameters.

**Figure 2.**
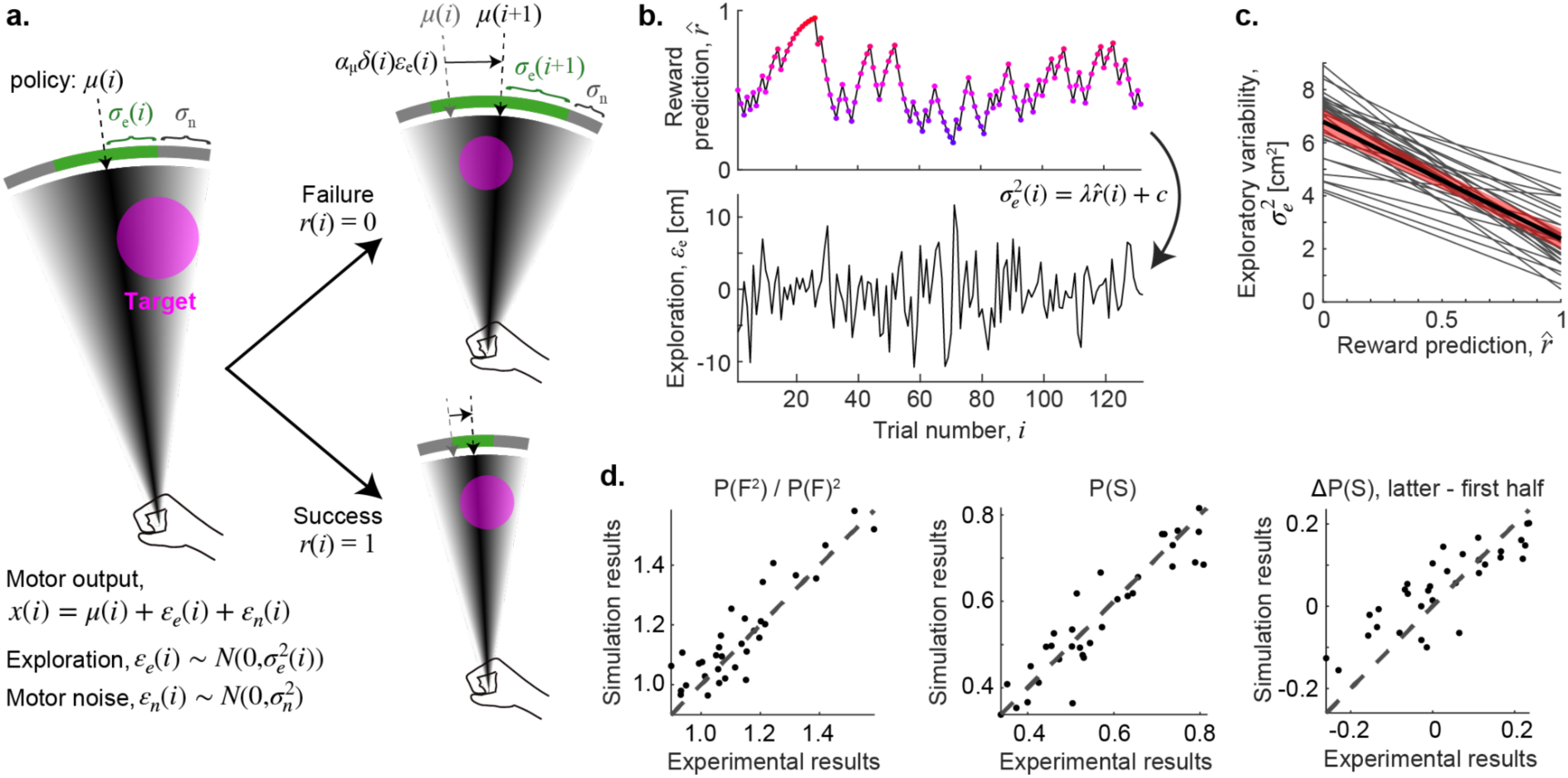
Reward prediction influences motor variability through exploratory action, leading to successive successes and failures. (a) Schema of a reinforcement learning model that updates policy direction *µ* and exploratory variability *σ_e_* based on trial outcomes. Motor noise *σ_n_* and learning rate *α_µ_* are constant free parameters. (b) In the model, exploratory variability has a negative linear relationship with reward prediction. The scatter plot shows how exploratory variability changes over trials with varying reward predictions. (c) Variability control function. Slope *λ* and intercept *c* were estimated for each participant using simulations with the reinforcement learning model. The red shaded area represents the 95% confidence interval. (d) Comparison of empirical results with simulation results. The model replicated (*left*) the emergence of successive failures, shown by the ratio of the probability of two successive failures P(F^2^) to the second power of the failure probability for the main examination P(F)^2^, (*middle*) the overall success rate, and (*right*) the difference in success rates between the first and latter halves of the experiment for each individual. The dotted lines in each figure represent perfect replication of the empirical results by the model-based simulations.

Four free parameters in the model (*α_µ_*, *σ_n_^2^*, *λ*, and *c*) were individually estimated to replicate three essential features of the experimental results: the ratio between the observed frequency of two SFs and their theoretical value, the overall success rate, and the difference in success rate between the first and second halves of the experiment. Simulation results using the computational model successfully replicated the experimental results with the estimated parameters, where the values (mean ± 95%CI) were *α_µ_* = 0.27 ± 0.02, *σ_n_^2^* = 2.76 ± 0.45, *λ* = −4.41 ± 0.54, and *c* = 6.79 ± 0.45. The model accurately predicted the occurrence of SFs (Fig. 2d left; R^2^ = 0.802), the overall success rate (Fig. 2d middle; R^2^ = 0.848), and the difference in the success rate between the first and second halves of the experiment (Fig. 2d right; R^2^ = 0.709).

### No association between motor performance improvements and the emergence of SFs

Given that our results showed that failure in one trial leads to failure in subsequent trials due to motor exploration, we posed the following question: Are there any links between SFs and motor learning outcomes? To investigate this, we classified the 33 participants into two groups: learners, who exhibited higher success rates in the second half of the main examination compared to the first half, and non-learners, who exhibited lower success rates in the second half compared to the first half. Thus, we obtained 17 learners, 14 non-learners, and two participants whose success rates remained unchanged between the first and second halves (Fig. 3a).

**Figure 3.**
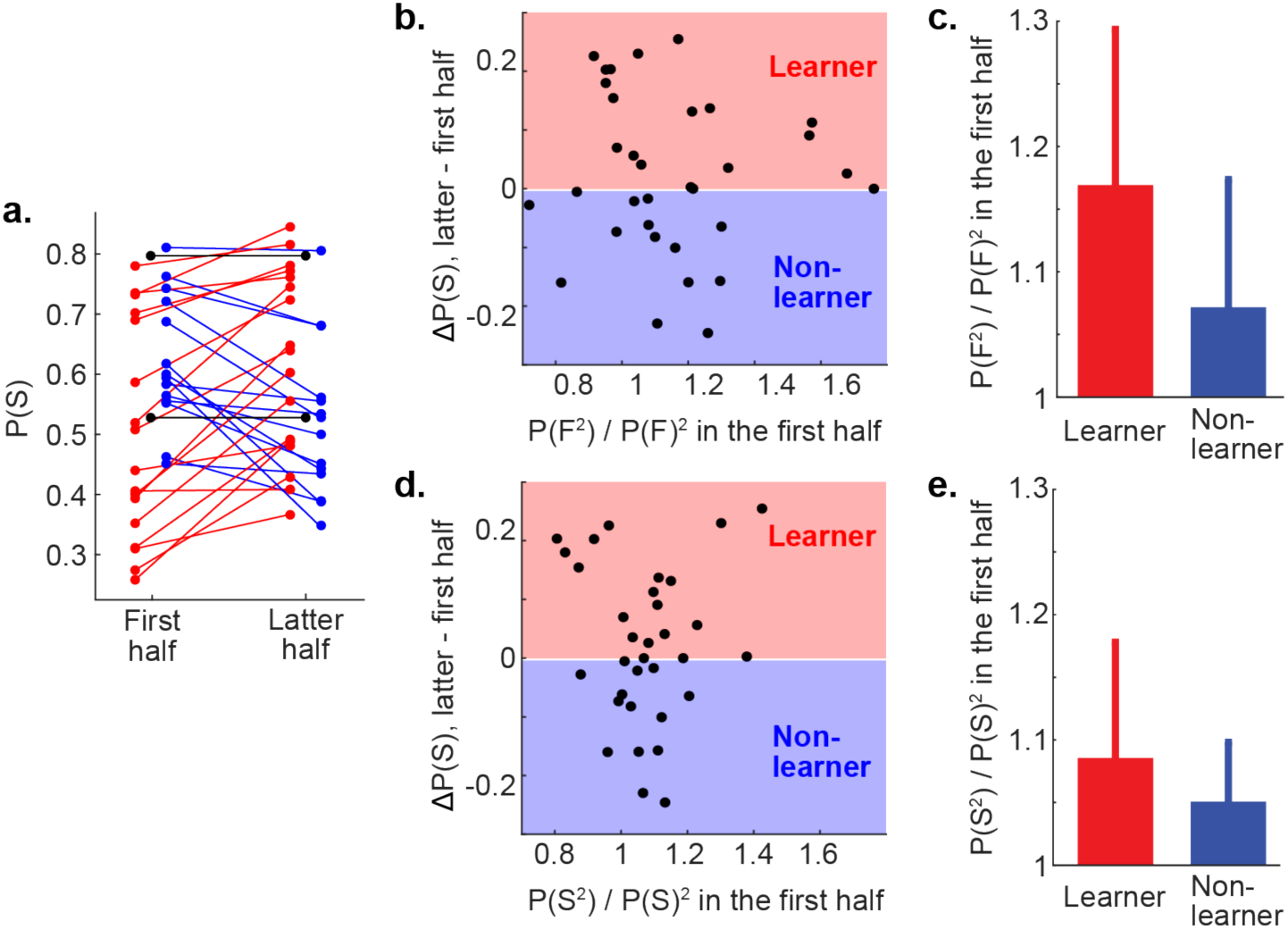
Lack of association between motor performance improvements and the emergence of successive failures or successes. (a) Success rates in the first and latter halves of the main examination. Participants were categorized as learners (red) and non-learners (blue) based on changes in success rates between the two epochs. Two participants (black) showed no change in success rate. (b) Scatter plot showing no correlation between motor performance improvement and the emergence of two successive failures during the first half of the examination. (c) Comparison of the emergence of two successive failures in the first half between learners and non-learners (mean ± 95% confidence intervals). (d) Scatter plot showing no correlation between motor performance improvement and the emergence of two successive successes during the first half of the examination. (e) Comparison of the emergence of two successive successes in the first half between learners and non-learners (mean ± 95% confidence intervals).

Our results suggest no association between motor performance improvements and the emergence of SFs or successes. No significant correlation existed between the change in the success rate from the first to the second halves of the main examination and the ratio of observed to theoretical frequencies of two SFs in the first half (Fig. 3b; r(31) = −0.069, p = 0.70). Similarly, no significant difference was found in the probability of SFs in the first half between learners (mean ± 95%CI: 1.170 ± 0.124) and non-learners (mean ± 95%CI: 1.072 ± 0.102) (Fig. 3c; t(29) = 1.26, p = 0.22). For successive successes, no correlation was observed between the change in success rate and the ratio of observed to theoretical frequencies in the first half (Fig. 3d; r(31) = −0.011, p = 0.95). Furthermore, no difference existed in the probability of successive successes between learners (mean ± 95%CI: 1.086 ± 0.092) and non-learners (mean ± 95%CI: 1.054 ± 0.048) (Fig. 3e; t(29) = 0.67, p = 0.51).

We next explored the association between motor performance improvements (measured by the difference in success rate between the first and second halves of the main examination) and several model parameters that affect motor outputs in different ways. Correlational analyses revealed a statistically significant relationship between motor performance improvement and exploratory variability when expecting maximal reward (Fig. S2a; r(31) = −0.41, p = 0.017). In other words, reduced exploratory variability, coupled with strong confidence in task success, was associated with improved motor learning. However, performance improvement was not correlated with exploratory variability when reward prediction equaled zero (Fig. S2b; r(31) = 0.093, p = 0.61), the gain of the exploratory variability control function *λ* (Fig. S2c; r(31) = −0.31, p = 0.082), or the degree of motor noise *σ_n_^2^* (Fig. S2d; r(31) = 0.16, p = 0.36).

### Emergence of successive successes and failures was task dependent

To further investigate the emergence of successive successes and failures, we examined whether these patterns were specific to tasks with identical targets or if they extended to conditions with multiple variable targets. This question stems from the idea that environmental variability, such as the presence of multiple targets, may influence exploratory variability shaped by recent reward history (Dwahale et al., 2019). In other words, do changes in the task environment disrupt or maintain the streaks of successes and failures observed with a single target? To address this question, a follow-up experiment was conducted with seven targets uniformly distributed over a ±90° range during the main examination (Fig. 4a). As in Experiment 1, 11 healthy young adults first underwent baseline testing in an environment with a single target to estimate the target size for 50% success probability. In the main examination, one of seven targets of equal size was randomly presented for each trial.

**Figure 4.**
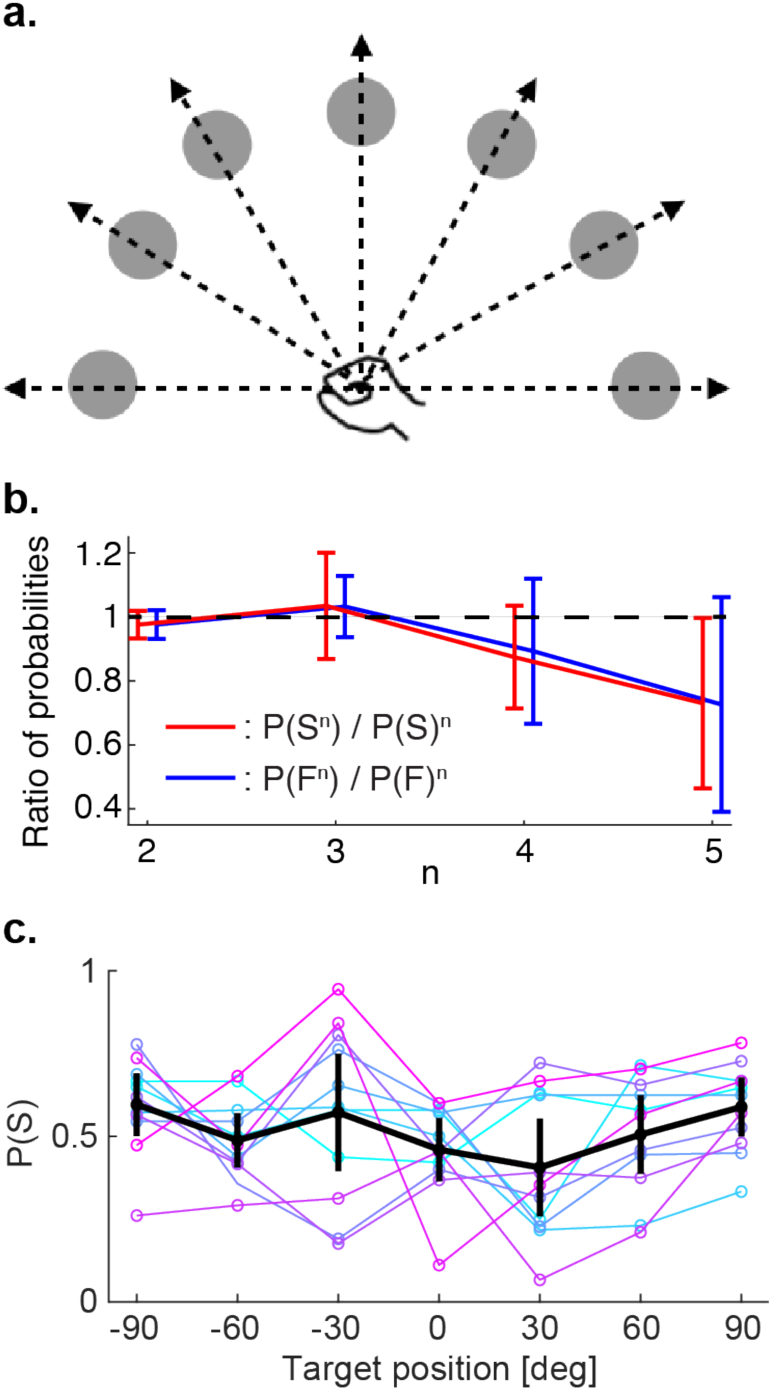
Emergence of success and failure streaks were task dependent. (a) A follow-up experiment was conducted with the same experimental procedure as in Experiment 1, except one of seven targets, uniformly distributed within a ±90° range, was randomly shown at each trial during the main examination. (b) Assessment of successive successes and failures. The red line represents the ratio of the probability of *n* successive successes P(S)^n^ and the *n*-th power of the success probability for the main examination P(S)^n^. The blue line represents the ratio of the probability of *n* successive failures P(F)^n^ and the *n*-th power of the failure probability for the main examination P(F)^n^. The probabilities of occurrence of *n* successive successes and failures did not differ from chance levels. Data are represented as the mean ± 95% confidence interval. (c) Success rates of reaching movements varied across target positions. Data are represented as the mean ± 95% confidence interval.

The results demonstrated that the observed frequencies of two successive successes and failures were not statistically different from their theoretical values, which are the square of the success/failure probability for the entire main examination (Fig. 4b). The exact values of the ratio between the observed frequency of two successive successes and failures and their theoretical values were 0.964 ± 0.043 and 0.961 ± 0.046 (mean ± 95%CI), respectively. We also investigated whether the success rate differed depending on the target location. A one-way repeated measures ANOVA showed a significant main effect of the target position on the success rate (F(6, 60) = 2.58, p = 0.027). However, post hoc pairwise comparisons with Bonferroni correction did not yield significant differences in the success rates among target positions.

## Discussion

In this study, we examined the occurrence of successive successes and failures in a reinforcement-based motor task and explored the mechanisms underlying these phenomena using a computational model. Our results showed that successive successes and failures occurred more frequently than would be expected by chance, suggesting a patterned behavior in motor performance that deviates from random variation. Specifically, the observed probability of two successive successes was 1.086 times higher than the theoretical probability, and the probability of two SFs was 1.143 times higher. A computational model, in which MV comprised constant motor noise and exploratory variability regulated by recent reward history (where variability increased following failure), successfully explained the emergence of SFs. These findings indicate that binary success/failure feedback in motor tasks induces performance streaks, probably caused by neural processes that regulate motor exploration and exploitation.

Our findings align with the framework of reinforcement learning, which posits that failure promotes exploration by increasing MV, whereas success leads to exploitation by reducing variability (Barto et al., 2017). This pattern has been consistently observed across various motor learning studies, including arm-reaching tasks and visuomotor adaptation, where greater variability typically follows unsuccessful trials (Cashaback et al., 2019; Chen et al., 2017; Pekny et al., 2015; Sidarta et al., 2018; Uehara et al., 2019; van der Kooij et al., 2018; van Mastrigt et al. 2020). Several models of reinforcement motor learning, such as those proposed by Therrien et al. (2016, 2018), Cashaback et al. (2019), Dhawale et al. (2019), and van Mastrigt et al. (2020), share the common feature that exploration increases following failure rather than success. However, recent critiques have pointed out that performance-dependent feedback can cause a correlation between motor noise and exploration, particularly in successful trials, leading to biased estimates of exploration (van Mastrigt et al., 2021). Additionally, estimating exploration based on trial-to-trial changes can lead to an underestimation of actual exploration, as the reference trial itself may contain exploration (van Mastrigt et al., 2021). Despite these limitations, our computational modeling successfully explained how the modulation of MV based on reward history leads to the emergence of performance streaks.

Interestingly, our results did not show a significant relationship between the likelihood of SFs and motor learning outcomes, as measured by changes in performance between the first and second halves of the experiment. While motor exploration has been emphasized as a key process in reinforcement motor learning, this lack of association indicates that SFs alone may not directly contribute to optimal learning strategies. Instead, they may represent exploratory actions that sample the motor space without consistently translating into performance improvements within the timeframe of the current task. This finding highlights the complexity of the interplay between exploration and exploitation, where exploratory variability might be necessary but insufficient for optimizing performance outcomes in all contexts. Factors such as task difficulty and individual differences in cognitive control and motivation may play more dominant roles in shaping motor learning processes (Sanli et al., 2015; Anderson et al., 2021). Notably, maintaining an optimal balance between challenge and success, such as an error rate of approximately 32%, enhances learning by maximizing engagement and error sensitivity (Al-Fawakhiri et al., 2023; Bengtsson et al., 2009).

Our experiments with seven targets revealed that SFs did not occur, which may be partially explained by the contextual interference effect. Contextual interference, a well-documented phenomenon in motor learning, suggests that high interference (e.g., random practice involving varied tasks) promotes learning by enhancing memory reconsolidation and the integration of task-relevant information (Shea & Morgan, 1979). Specifically, the reconstruction hypothesis emphasizes the role of variability during memory reconsolidation in improving motor skill performance (Borges et al., 2023), while the elaborative processing hypothesis posits that random practice facilitates intertask comparisons and task-relevant elaboration (Lin et al., 2023). These mechanisms may have contributed to disrupting the conditions for SFs in the seven-target condition. However, the random presentation of seven targets in our study involved variations of the same motor task rather than distinct tasks, making it less clear whether the traditional contextual interference theory fully applies. Beyond contextual interference, temporal dynamics in reinforcement learning may provide complementary explanations. The separation between targets in the seven-target condition could lead to forgetting of learned values in reinforcement learning (Kato & Morita, 2016), potentially dampening exploratory variability to baseline levels, as participants could not act immediately on errors related to the same target. This interplay between task variability and memory processes warrants further investigation to fully elucidate the mechanisms underlying these findings.

Although studies directly linking successive successes and failures to specific neural activity are lacking, evidence from motor skill acquisition and reinforcement learning research suggests that the activity of several cortical regions is involved in processing performance outcomes. An fMRI study by Rubia et al. (2007) indicated that failure trials activate the mesial prefrontal cortex as well as the anterior and posterior cingulate gyri. Similarly, Zhao et al. (2016) demonstrated that the number of failures experienced during motor tasks is associated with activation in multiple brain regions, including the frontal, parietal, and temporal cortices, as well as the limbic system and striatum. Notably, changes in the anterior cingulate cortex, posterior cingulate cortex, and temporoparietal junction have been observed in response to repeated failures, suggesting that these regions play a critical role in processing error feedback and adapting motor strategies (Zhao et al., 2016). The fMRI study showed that locus coeruleus-norepinephrine (LC-NE) engagement in response to task-related salient events was associated with differential reaction time effects (Ludwig et al., 2024). Additionally, the LC-NE system, which plays a crucial role in regulating attention and behavior adaptation through reinforcement learning (Su et al., 2022), could be involved in the occurrence of serial successes and failures. Measuring pupil diameter, which correlates with LC-NE activity and serves as a noninvasive marker of neuromodulatory changes related to reinforcement learning and decision-making uncertainty (Gilzenrat et al., 2012; Geng et al., 2015), could offer insights into the neurophysiological basis of performance streaks in motor tasks.

Overall, the present study highlights the critical link between SFs and motor exploration, suggesting a mechanism behind individual differences in motor performance. While errors are often considered a necessary part of motor skill improvement rather than obstacles to be avoided (Seidler et al., 2013), the functional significance of SFs remains unclear. Our findings demonstrate that motor exploration, regulated by recent performance outcomes, can lead to performance streaks where SFs emerge as part of the natural trial-and-error process rather than purely negative outcomes. This suggests that failure streaks represent active exploration rather than events that should always be suppressed. However, the long-term effects of suppressing or allowing such failures on motor performance remain uncertain. Future research should investigate whether suppressing SFs hampers or enhances skill acquisition to provide further insights into how MV fluctuations contribute to optimal motor learning.

## Limitations of the study

One limitation of this study is the lack of neurophysiological data. While our findings point to a link between SFs and motor exploration, we did not measure neural activity directly, which limits our ability to draw concrete conclusions about the underlying neurophysiological mechanisms. Future research using techniques such as fMRI, EEG, or pupil diameter measurements could provide more insight into how neural systems support or modulate these failure streaks. Additionally, our study did not investigate the effects of inhibiting SFs. The relationship between the suppression of failure streaks and motor performance remains unexplored, leaving open questions about whether such inhibition would enhance or impair learning. Future studies should examine how modifying the occurrence of SFs might influence the overall learning process.

## Materials and methods

### Participants

We recruited 49 healthy adults for this study. All participants were right-handed, had normal or corrected-to-normal vision, had no history of neurological disease, and had not participated in any experiments conducted by our group in the last 6 months. Five participants were excluded from the statistical analysis because their data met the exclusion criteria (see *Experimental procedure* for details). Consequently, data from 44 adults (31 males and 12 females; age 21.9 ± 2.6 years, mean ± SD) were analyzed. The study was conducted in accordance with the Declaration of Helsinki. The experimental procedures were approved by the ethics committee of the University of Tokyo (approval number: 18-203). Written informed consent was obtained from all participants prior to the experiments.

### Apparatus and general experimental task

Participants sat on a chair and grasped the handle of a robotic manipulandum (KINARM End-Point Lab, Bkin Technologies, Ontario, Canada) with their right hand. They moved the handle horizontally from a home position toward a target displayed on a monitor placed above their arm. Although participants could not see their own arm directly, they could see the handle position represented by a gray circle (diameter = 1 cm) on the monitor. The upper body was secured to the chair with straps to minimize the movement of other body parts. The handle position was sampled at 1,000 Hz (BNC-2090A, National Instruments, Texas, USA) and low-pass filtered using a second-order Butterworth filter with a cutoff frequency of 10 Hz. Handle velocity was obtained through numerical differentiation of the handle position data.

The start position for the reaching movements was set at 15 cm from the edge of the monitor (approximately 20–30 cm from the chest). The target was located 10 cm away from the start position. Both the start and target positions were displayed as circles. The target radius was adjusted based on the participants’ performance, with the radius at the beginning of the baseline examination set at 8 cm. After the participants maintained their right hand at the start position for 1–1.5 s, a white target appeared on the monitor. Following an additional waiting time of 1.25 ± 0.25 s, the target color changed to magenta, indicating that participants should reach toward the target. Right after the start of the movement, the visual feedback of the handle position disappeared. Once participants stopped their hand movements, they received binary feedback of success or failure, defined by whether the hand path overlapped with the target. However, if the peak velocity of their hand movement was slower than 40 cm/s, a warning message, “Slow,” was displayed on the monitor and no binary feedback was provided. Participants were instructed to make their movements achieve “Success” and avoid showing “Slow” as much as possible. After participants did not move their right hand for 1.5 s, the robot returned the handle to the start position. Participants were asked to hold the handle continuously throughout the trials.

### Experimental procedure

#### Experiment 1: Single reaching target

First, the participants familiarized themselves with the motor task by performing 20–30 reaching movements to the target in front of the start position. During this period, the target radius was fixed at 8 cm, and either “Success,” “Failure,” or “Slow” was shown on the monitor as performance feedback. The baseline examination was then conducted for approximately 10 min (59–78 trials) to estimate the target size for a 50% success rate. During the baseline examination, the target radius decreased by 1 cm following success, increased by 1 cm following failure, and remained unchanged if the movement was slower than 40 cm/s (Fig. 1b). After the baseline examination experiment, we individually calculated the success rate for each target radius and fitted a sigmoidal curve to estimate the target radius for a 50% success rate, which was then used for the main examination. The main examination lasted 20–40 min (124– 302 trials). The target was always shown at the front of the start position with a constant radius. Data were originally obtained from 38 participants, but we omitted data from five individuals whose estimated target radius for the main examination was over 8 cm.

#### Experiment 2: Multiple reaching targets

The experimental procedure of Experiment 2 was almost identical to that of Experiment 1, except for the number of targets during the main examination. Seven targets were located at 0° (in front of the start position), ±30°, ±60°, and ±90°. The same radius was used for all seven targets. The target position was pseudorandomly shifted in every trial. The main examination lasted 20–25 min (136–164 trials). Data were obtained from 11 participants, and none of the participants met the exclusion criteria.

### Computational simulations

#### Reinforcement learning model

We modeled a motor task in a one-dimensional continuous motor space, referring to a previous study (Dhawale et al., 2019). Movement direction (*x*) was drawn from a stochastic Gaussian policy with mean *μ* and variance comprising two components – exploratory variability ε_*e*_ ~ *N*(0, σ^2^_*e*_) and motor noise ε_*n*_ ∼ *N*(0, σ^2^_*n*_) (Fig. 2a):

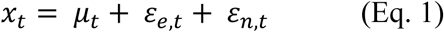

where *t* is the trial number in a simulated session and *N*(0, σ^2^) references a normal distribution with mean 0 and variance σ^2^. Binary reward *r* ∈ {0,1} was available only within a circumscribed width of angle w centered at 0 degrees (Fig. 5a, top and middle). The agent updated the mean movement policy in the direction of the rewarding angle using an on-policy learning rule:

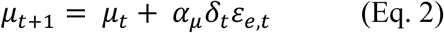

**Figure 5.**
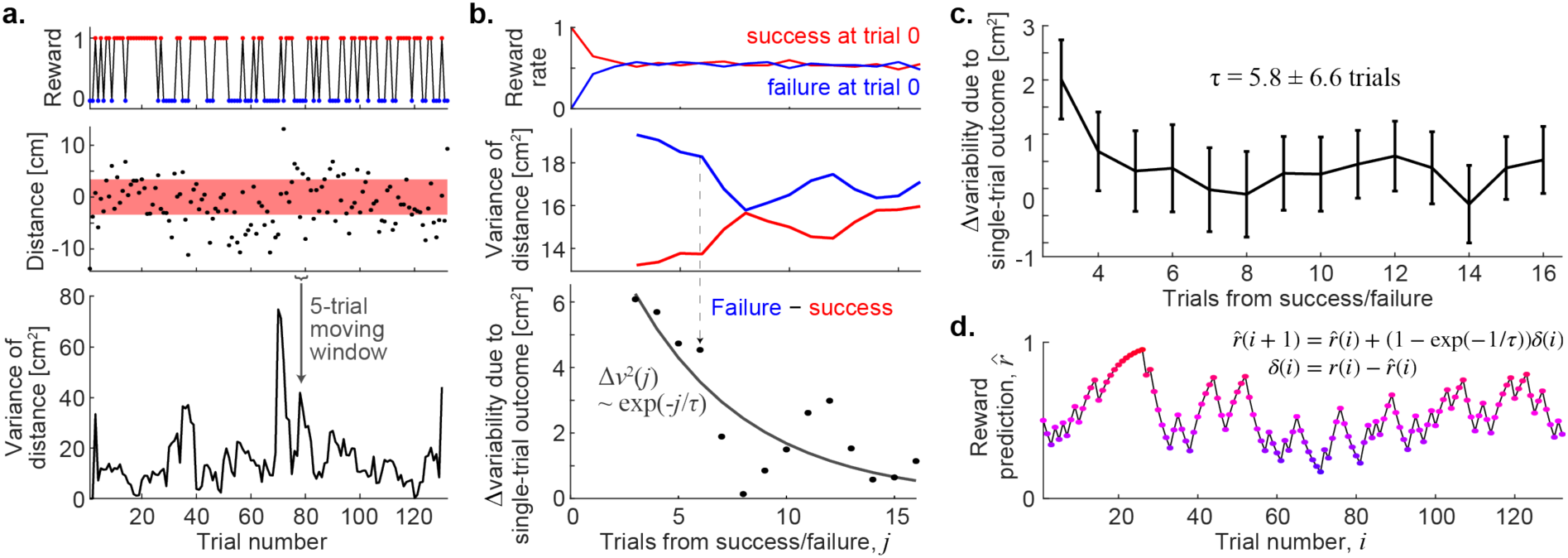
Reward prediction was computed based on recent reward history and movement variability. (a) *Top*: Reward values (success or failure) of a representative participant. *Middle*: Minimum distance from the target center during a single trial of reaching movement. The red shaded area indicates the target size. *Bottom*: Variance in reaching movements. (b) *Top*: Performance-triggered average reward rate for the same participant, as shown in Figure S3a top. *Middle*: Average motor variability conditioned on success (red) and failure (blue) in a single trial (at trial 0). *Bottom*: The decay rate of single-trial variability effect (*τ*) was estimated using exponential fitting to the difference between average motor variability in response to single-trial success and failure. Variabilities at trials 1 and 2, which were significantly influenced by performance at trial 0, were excluded to minimize the confounding influence of performance at trial 0 on the regulation of future motor variability. (c) Difference between motor variability in response to single-trial performances, averaged across 33 participants. Data are represented as mean ± SEM. (d) Reward predictions for the same participant. These values were updated using a reward prediction error *δ* multiplying by the decay rate of single-trial variability effects, as shown in Figure 5a (bottom).

where δ_*t*_ = *r_t_* − *r̄*_*t*_ represents the reward prediction error and α_μ_ is the learning rate for the policy mean. The reward prediction (*r̄*) was estimated from the sequence of rewards using the following learning rule:

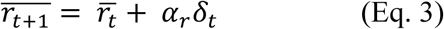

where α_*r*_ is a parameter for updating the reward prediction. We also implemented reward-dependent modulation of variability by designating a variability control function to set levels of exploratory variability as a function of reward prediction (Fig. 2c):

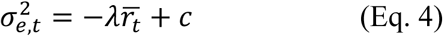

Unlike the previous study (Dhawale et al., 2019), our Experiment 1 did not include environmental variability. Thus, we simplified their model while retaining the essential feature: the negative relationship between reward rate and exploratory variability. This simplification allowed us to focus on the core interaction between reward history and MV.

#### Analysis of MV modulation

To perform simulations, we analyzed how the outcome of a single trial (i.e., success or failure) modulated MV in subsequent trials (Fig. 5). Initially, we computed the variance in movement direction, which is the minimum distance from the center of the target in a single trial, using a 5-trial moving window (Fig. 5a). We then assessed the differences between reinforcement-triggered averages of movement direction variability following the success and failure trials for each participant to evaluate the effect of a single-trial outcome on MV (Fig. 5b). The results showed that recent performance played a significant role in modulating future MV—variability increased after failure trials and decreased after success. Furthermore, the effect of trial outcomes on variability decayed exponentially, with a participant-averaged time constant of 5.8 ± 6.6 trials (mean ± 95%CI) (Fig. 5c). The update rate for reward prediction, denoted as α_0_, was calculated for each participant based on their individual time constant *τ*:

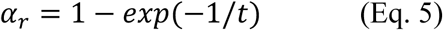

This equation allowed us to compute the reward prediction (Fig. 5d) by convolving past trial outcomes with a reward rate filter (i.e., essentially a weighted average of past outcomes) derived from the exponential decay of the single-trial variability effect.

#### Simulations

We simulated a one-dimensional motor task for individual participants using the reinforcement learning model described earlier. Session-specific parameters, including the starting position relative to the center of the target (μ_0_), the width of the success angle (w), and the number of trials, were derived from each participant’s dataset. The update rate for reward prediction, denoted as α_*r*_, was also calculated from the analysis of individual participant data. To optimize the learning rate for the policy mean α_μ_, the magnitude of motor noise σ^2^_*n*_, and the two parameters governing the variability control function (λ and c) for each participant, we simulated task performance across a range of values (α_μ_: 0.20–0.40, σ^2^_*n*_: 0.50–6.50, λ: 0.50– 7.50, c: 3.50–9.50). The optimal parameter values were determined by minimizing the sum of the mean-squared errors between simulated outcomes and the experimentally observed results during the main examination: the overall success rate and the difference in success rate between the first and second halves of the main examination.

### Statistical analyses

We confirmed that all values used in the statistical analyses met the criteria for normal distribution using the Lilliefors test. Therefore, correlation analyses were conducted by calculating Person’s correlation coefficients, and pairwise comparisons were performed using either paired t-tests or two-sample t-tests. One-way repeated measures ANOVA was conducted without Greenhouse–Geisser correction, as Mauchly’s sphericity test confirmed that the assumption of sphericity was not violated. The type I error threshold was set at 0.05 for all statistical analyses.

### Software

Data analysis, computational simulations, and statistical analyses were performed using MATLAB 2020b (Mathworks, Massachusetts, USA).

## Acknowledgements

This work was supported by JSPS KAKENHI to MT (20K19593) and to DN (21H04860).

## Author contributions

Conceptualization: RS, MT, DN; Data curation: RS, TO; Formal analysis: RS, MT, MK; Funding acquisition: MT, DN; Investigation: RS, TO; Methodology: RS, MT, MK; Supervision: MT, DN; Visualization: RS, MT; Writing – original draft: RS, MT. Writing – review & editing: RS, MT, MK, DN.

## Declaration of generative AI and AI-assisted technologies in the writing process

During the preparation of this work the authors used ChatGPT in order to correct grammatical errors and to improve the readability. After using this tool, the authors reviewed and edited the content as needed and take full responsibility for the content of the published article.

## Data and code availability

Data and code to generate all figures associated with the results can be found at https://osf.io/r654y/. The other data (e.g., raw data) will be provided upon request to the lead contact, Mitsuaki Takemi.

## Supplementary information

**Figure S1.**
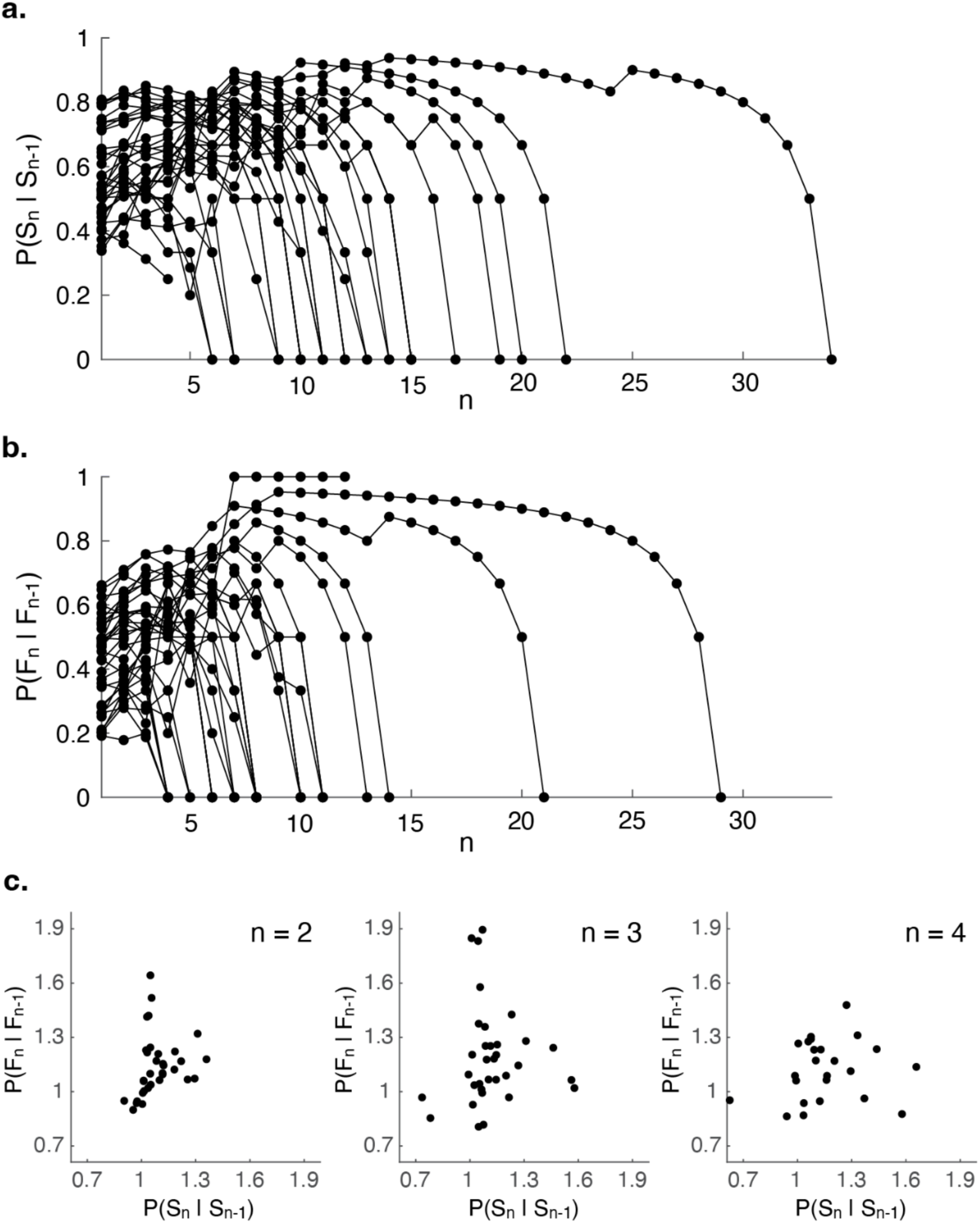
Probabilities of successive successes and failures as a function of trial number. (a) Probability of achieving success in the current trial *n* given *n* − 1 consecutive prior successes P(S_n_ | S_n-1_) for each participant. (b) Probability of failure in the current trial *n* given *n* − 1 consecutive prior failures P(F_n_ | F_n-1_) for each participant. Each line represents data from an individual participant. (c) The relationship between the probabilities of successive successes P(S_n_ | S_n-1_) and successive failures P(F_n_ | F_n-1_) for each participant at different trial lengths (n = 2, 3, 4). Each point represents data from an individual participant.

**Figure S2.**
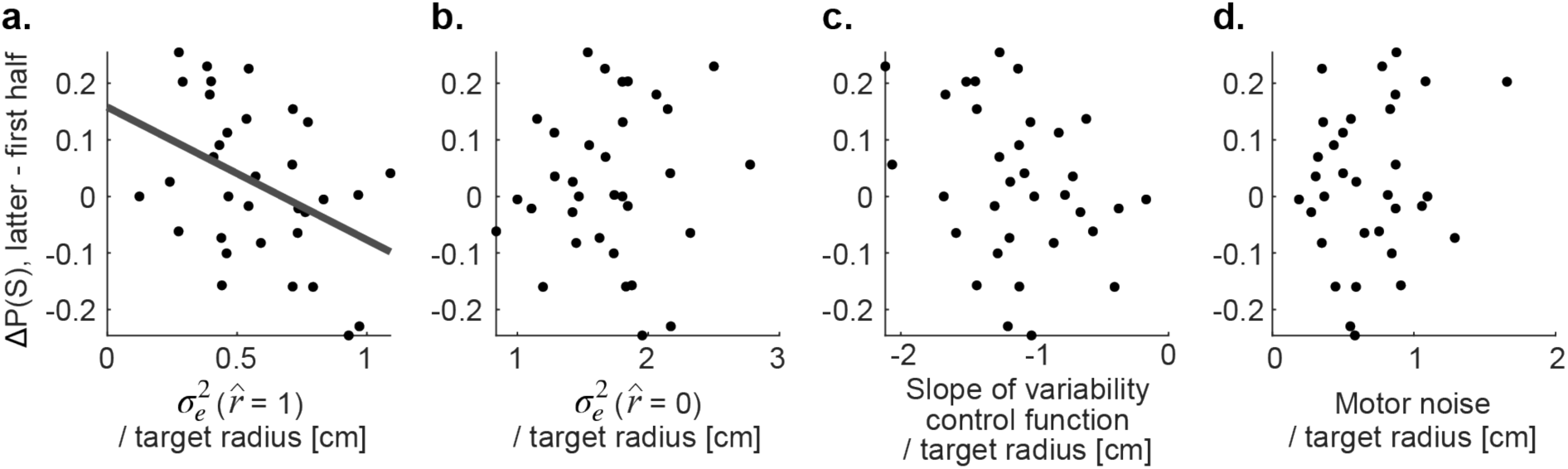
Associations between changes in success rates during the main examination and model parameters. Each panel illustrates the correlation between specific model parameters and the difference in success rates between the latter and first halves of the examination (ΔP(S)). Significant correlations are indicated with least-squares regression lines. (a) Relationship between exploratory variability under the expectation of maximal reward and ΔP(S). (b) Relationship between exploratory variability when reward prediction equals zero and ΔP(S). (c) Relationship between the slope of the variability control function and ΔP(S). (d) Relationship between motor noise and ΔP(S).

